# Expansion Strategy-Driven Micron-Level Resolution Mass Spectrometry Imaging of Lipids in Mouse Brain Tissue

**DOI:** 10.1101/2023.08.28.555097

**Authors:** Yik Ling Winnie Hung, Chengyi Xie, Jianing Wang, Xin Diao, Ruxin Li, Xiaoxiao Wang, Shulan Qiu, Jiacheng Fang, Zongwei Cai

## Abstract

A novel method for enhanced resolution, termed expansion mass spectrometry imaging (Ex-MSI), has been developed for lipid mass spectrometry imaging, utilizing existing commercially available mass spectrometers without necessitating modifications. This approach involves embedding tissue sections in a swellable polyelectrolyte gel, with the target biomolecules indirectly anchored to the gel network. By employing matrix-assisted laser desorption ionization mass spectrometry imaging (MALDI-MSI), the method has realized an enhancement in spatial resolution that surpasses the conventional resolution limits of commercial instruments by approximately 4.5 folds. This enhancement permits the detailed visualization of intricate structures within the mouse brain at a subcellular level, with a lateral resolution nearing 1 μm. As a physical technique for achieving resolution beyond standard capabilities, this approach is readily adaptable and presents a powerful tool for high-definition imaging in biological research.

## Introduction

In recent decades, the need for high spatial resolution imaging has grown significantly across various disciplines in biological research^[1-3]^. This technology has become a pivotal tool in not only identifying chemical components within cells and tissues but also elucidating intricate cellular structures, functions, and interactions. In the context of disease, high spatial resolution imaging is invaluable for detecting biomarkers, thus playing a crucial role in the prevention, diagnosis, treatment, and monitoring of various conditions. Beyond biomedicine, it also fosters progress in fundamental research, aiding in the understanding of biological processes at the molecular and subcellular levels. Such insights are integral to discoveries in fields ranging from cell biology to neuroscience^[4]^.

Confocal or super-resolution fluorescence microscopy is a widely used technique for high spatial-resolution molecular imaging. However, this approach has inherent limitations^[5, 6]^. Its reliance on fluorescent dyes and antibody recognition can introduce complexities, particularly when multiple dyes and antibodies are needed to investigate different molecular species within a single experiment. This multiplicity often leads to signal interference and decreased throughput. While these microscopy-based methods are efficient in visualizing the spatial localization of proteins, they face challenges in imaging small molecules with high molecular specificity. This drawback is primarily due to the difficulties in creating specific recognition tags for small molecules, which is crucial for accurate molecular imaging.

Mass spectrometry imaging (MSI) has emerged to address these challenges^[7-9]^. MSI enables the direct imaging of numerous chemical species in a single experiment. This includes the ability to detect both large biomolecules, such as proteins and peptides, and smaller biomolecules, like lipids and metabolites. Unlike fluorescence microscopy, MSI does not rely on complex labeling processes or the use of exogenous dyes. This attribute enhances MSI’s versatility and effectiveness in studying cellular structures and functions. Notably, secondary ion mass spectrometry (SIMS) achieves high spatial resolution, capable of reaching sub-micrometer resolution. However, the primary limitation of SIMS is severe ion fragmentation during analysis, which can diminish its sensitivity and potentially impact the accuracy of molecular identification^[10]^. Alternative techniques such as multiplexed ion beam imaging (MIBI) and imaging mass cytometry (IMC) demonstrate remarkable resolution capabilities at the sub-micrometer level for high multiplex protein imaging^[11-15]^. However, a significant constraint is the requirement for expensive immuno-tags, which can elevate the overall cost of the procedures. Additionally, these techniques often face limitations in throughput, generally being capped at fewer than 100 per experiment. Furthermore, MIBI and IMC are methods aimed at targeted spatial proteomics and are not suitable for analyzing metabolites and lipids.

Matrix-assisted laser desorption ionization mass spectrometry imaging (MALDI-MSI) has emerged as a promising method in the pursuit of high spatial resolution molecular imaging^[10, 16-27]^. This technique has the distinct advantage of retaining various intact biomolecules without the need for labeling, making it particularly appealing for studying complex biological structures. Among the available soft ionization techniques, MALDI-MSI offers the highest spatial resolution, with commercial settings achieving 5-μm resolution. Recent innovations in mass spectrometry imaging techniques, such as scanning microprobe MALDI (SMALDI), have significantly extended the boundaries of spatial resolution. These advancements have enabled SMALDI to achieve resolutions as fine as 1.4 μm^[16]^. The transmission mode optics in mass spectrometry imaging (tMALDI), has led to substantial advancements in spatial resolution. Utilizing this technique, certain experiments have successfully achieved resolutions of 1.2 μm and even finer resolutions down to 0.6 μm through oversampling ^[18]^. However, these remarkable achievements often come with challenges. They may require customized instrumentation, sophisticated optics design, and intricate sample preparation. These complexities can hinder the broad adoption of these methods, posing obstacles to their widespread commercialization.

In response to these challenges, we have developed Expansion Mass Spectrometry Imaging (Ex-MSI), an innovative method that increases the spatial resolution of MALDI-MSI to the subcellular level without the need for modifications in laser optics. Inspired by expansion microscopy^[28-39]^, Ex-MSI physically expands the tissue slice embedded in an expandable hydrogel, achieving a ∼4.5-fold increase in spatial resolution without relying on the optics of mass spectrometer. This unique approach does not necessitate an ultra-fine matrix deposition method and can be readily adopted across different instruments. In this work, we demonstrate the effectiveness of lipid Ex-MSI across mouse brain with diverse structural properties at subcellular level resolution. We reveal the method’s capability to delineate fine and compact structures at a 1-μm resolution level. The Ex-MSI method holds significant promise for the broader application of high lateral spatial resolution MALDI-MSI in biological and biomedical disciplines, potentially accelerating medical research and drug development in the future.

## Results and Discussion

### The Expansion Mass Spectrometry Imaging Workflow

In this study, we utilized gelation formulations from expansion microscopy (ExM) ^[28, 29, 34, 36-39]^ and customized sample preparation methods to achieve ultra-high-resolution mass spectrometry imaging through effective physical expansion of the sample. Fig. 1 shows the schematic workflow of Ex-MSI. Fresh mouse brain was first fixed. The anchor molecule Acryloyl-X SE(AcX) was then linked to the proteins in tissue slices. Differing from traditional ExM, the tissue slice for MSI is required to lay flat and adhere uniformly to the gelation chamber’s bottom. During gelation, proteins were anchored to the gel network and lipids could be retained through electrostatic interactions between proteins and lipids in mild conditions. The tissue’s dense structure is disrupted during digestion to allow uniform expansion. Subsequently, the hydrogel is dried onto an ITO slide. To achieve high spatial resolution MSI, fine matrix droplets were deposited onto the hydrogel using pneumatic-assisted electrospray deposition. The mass spectrometry imaging parameters are optimized to enhance the signal-to-noise ratio of lipid peaks. After MS acquisition, the data was analyzed by SCiLS Lab software.

**Fig. 1:**
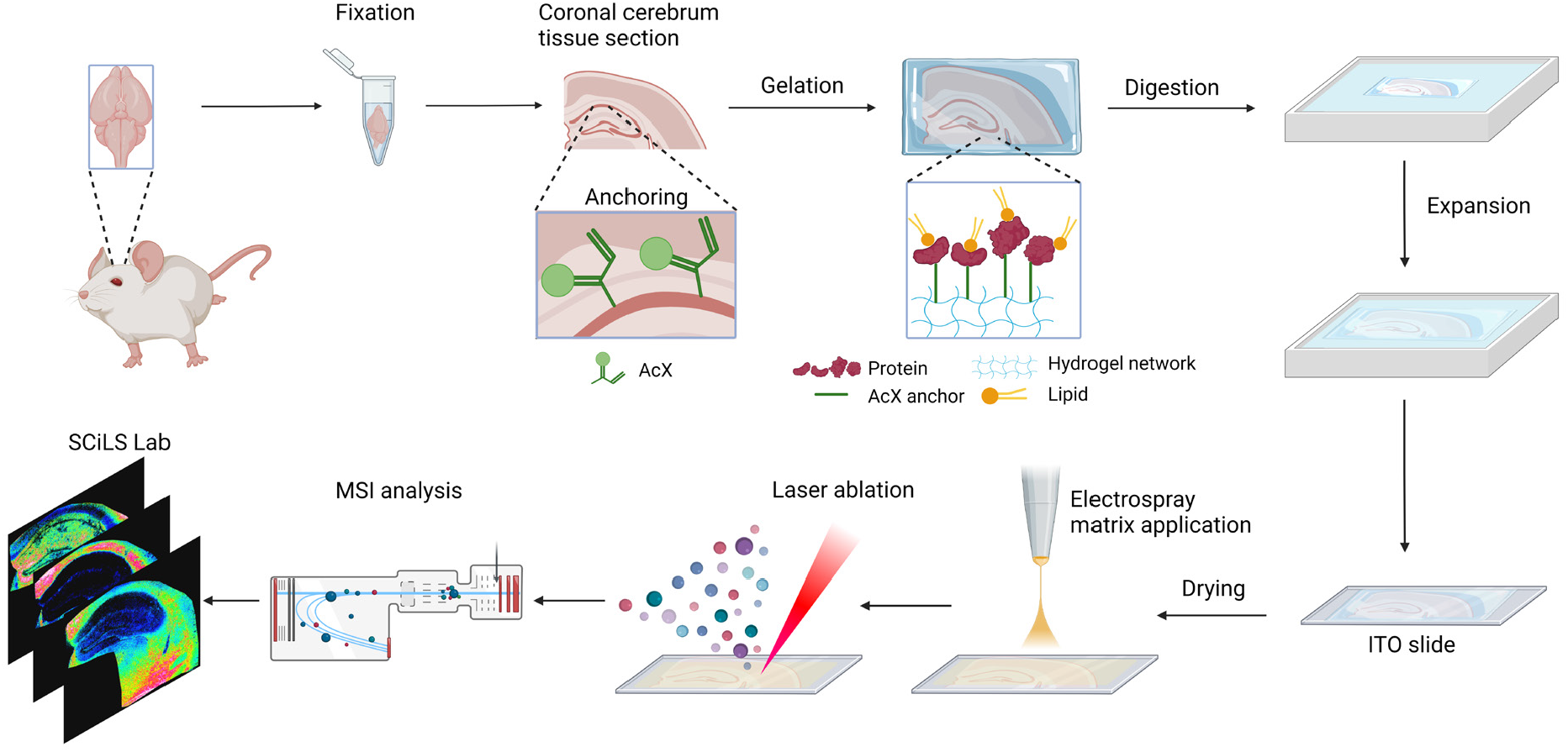
Schematic Overview of Ex-MSI Workflow. This figure provides a step-by-step illustration of the experimental procedures involved in Ex-MSI (Expansion Mass Spectrometry Imaging), from sample preparation to data acquisition and analysis. Created with BioRender.com.

After building the basic framework of the Ex-MSI workflow, the expansion factor and uniformity were determined. Concerning the expansion method, Suppl. Fig. 1a presents the mouse brain hippocampus region embedded in the gel before expansion, where the bottom width was 4.5 mm. After expansion, the bottom width became 26 mm (Suppl. Fig 1b) and it remained 23mm after drying (Suppl. Fig. 1c). The expansion factor could reach up to 5 folds (∼4.5 folds on average). In Suppl. Fig. 1d, the optical image obtained by bright-field microscopy and the MSI image exhibited on the left and right, respectively. The relative position and the shapes of the critical features in both images resemble each other. On the other hand, we performed MALDI-MSI for consecutive tissue slices to ascertain expansion uniformity. The MSI images of m/z 419.2 shows the outline of the tissue (Suppl. Fig. 1e). We separately performed with (right image) and without (left image) tissue expansion MSI for two adjacent tissue sections. The overlapped stacking comparison in Suppl. Fig. 1f showed that there was no obvious distortion in the shape of the tissue slices after overlaying the two MSI images together. The Ex-MSI method could achieve uniform tissue expansion without causing significant distortions or loss of fidelity.

To improve the efficiency and reliability of our method, we optimized the thickness of tissue sections in alignment with our gel formulation and the comprehensive sample preparation process. We observed that varying the thickness of tissue sections significantly impacts the ease of operation and the success rate. Thicker tissue sections tend to curl less, facilitating smoother flattening and ensuring more consistent adhesion of the tissue surface to the base of the entire gel. Suppl. Fig. 2a shows the signal intensity comparison of lipid Ex-MSI using 30, 40, and 50-μm tissue sections. Representative lipids, five phosphatidylcholines (PC) and five sphingomyelins (SM) were selected as listed in Suppl. Table 1 and their signal intensities in ten random ROIs across the coronal cerebrum tissue section were averaged. Among the ten lipids, 50-μm slices generally provided the highest signal intensities. As the tissue thickness increased from 30 to 50 μm, we observed a near doubling of signal intensities (Suppl. Fig. 2b). It’s not that thinner sections are incapable of generating equally strong signals, but rather, due to their mechanical properties and flatting behavior in the gel, the likelihood of achieving similar signal quality decreases. Thus, 50-μm slices would be used in the following experiments.

### Ex-MSI of lipids in mouse brain

To investigate the effect on imaging quality improvement after expansion, we imaged mouse brain regions using different spatial resolutions and the results were compared in Fig. 2. By increasing the laser rastering acquisition resolution from 50 to 5 μm, which was equivalent to the imaging resolution of 11 to 1.1 μm, the Ex-MSI shows significantly improved image resolution.

**Fig. 2:**
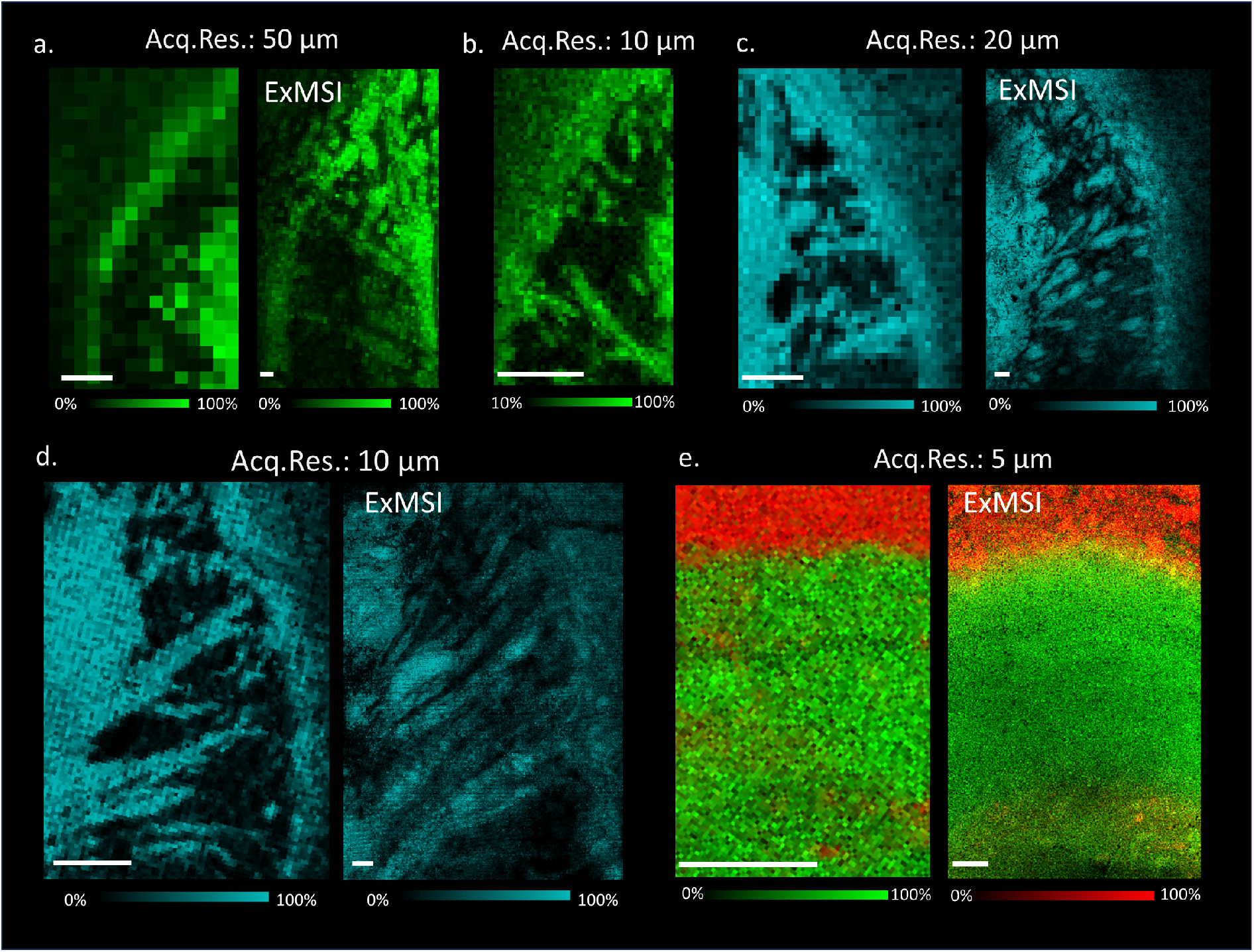
Ex-MSI of mouse cerebrum tissue slices. (a) Positive mode ion images of m/z 850.8, PC-O (36:4) [M+Na]^+^ in caudoputamen of the striatum region acquired with 50 μm acquisition resolution, resulting in an equivalent lateral spatial resolution of 11 μm for expanded group (right, labeled with ExMSI). (b) Positive mode ion image without expansion of m/z 850.8 in the same region acquired with 10 μm acquisition resolution. (c, d) Negative mode ion images of m/z 888.6, SHexCer (42:2;O2) [M-H]^-^ in caudoputamen of the striatum region acquired with 20 μm (c) and 10 μm (d) acquisition resolution, respectively, resulting in an equivalent lateral spatial resolution of 4.4 μm(c) and 2.2 μm (d) for expanded samples, respectively. (e) Negative mode ion images of m/z 888.6, SHexCer (42:2;O2) [M-H]^-^ (red), and m/z 718.5, PE (34:0) [M-H]^-^ (green) in alv and CA1 of the hippocampus region acquired with 5 μm acquisition resolution, resulting in an equivalent lateral spatial resolution of 1.1 μm for expanded sample. In (a, c, d, and e), the left and right panel displays the ion images between non-expanded and expanded group, respectively. Scale bar represents 200 μm.

To illustrate the effect of Ex-MSI, samples with and without expansion were prepared using consecutive coronal brain slices. The caudoputamen (CP) of the striatum region was imaged with 50 and 20 μm spatial resolution under positive mode with 2,5-Dihydroxybenzoic acid (DHB) as a matrix. In Fig. 2a, the ion images of m/z 850.8, [M+Na]^+^ ion of PC (O-36:4) are displayed. Without employing the expansion method (left panel), the characteristic fibrous morphology of CP is not discernible at a 50 μm acquisition resolution. However, when utilizing ExMSI (right panel), this morphology becomes visible even at a 50-μm acquisition resolution when expanded 4.5-fold, achieving an effective spatial resolution of 11 μm. For comparison, we directly imaged a slice of mouse brain cerebrum using a 10-μm laser raster spot produced ion images of similar quality (Fig. 2b), attesting to our method’s efficacy. When the CP was subjected to imaging at an acquisition resolution of 20 μm with expansion, translating to an equivalent image resolution of 4.4 μm, intricate structural details were clearly unveiled, as illustrated in Fig. 2c. The CP was furthermore imaged using 10 μm acquisition step width, achieved an equivalent resolution of 2.2 μm, the detailed structures emerged more clearly, as shown in Fig. 2d (right). By employing a more readily attainable 10-μm acquisition resolution, Ex-MSI allows for imaging capabilities that exceed the inherent resolution boundaries of the instrument. Moreover, this enhancement in resolution is independent of the instrument’s inherent capabilities, offering the potential for further improvement as the instrument’s resolution capabilities improve. After upgraded with the microGRID module, our instrument is capable of high-precision 5-μm acquisition without oversampling and misalignment. By conducting a 5-μm acquisition post-expansion, we achieved an impressive equivalent resolution of 1.1 μm. Fig. 2e presents mass spectrometry imaging of a localized area at the junction between the hippocampus and cortex, demonstrating the method’s capability to break through the 5-μm resolution limit and further reveal intricate details at micron-level resolution.

To further demonstrate Ex-MSI’s capability in capturing both complex morphology and intricate chemical compositions in brain tissue, we imaged a large region of a coronal brain slice at a 10-μm acquisition resolution in the negative ion mode. Fig. 3b shows an overlaid ion image at an equivalent 2.2-μm resolution for lipid species including PE (38:4) ([M-H]^-^, m/z 766.5), PE (40:6) ([M-H]^-^, m/z 790.5), and SHexCer (40:1;O2) ([M-H]^-^, m/z 862.6) via Ex-MSI, revealing their distinct distribution patterns within the brain. Figs. 3c-e provides enlarged ion images for various lipid classes, comparing the 2.2-μm equivalent resolution obtained through expansion (right or bottom panel) against the 10-μm native resolution (left or top panel). The regions of interest for these images are highlighted in Fig. 3b. Notably, the fibrous structure in the dorsal part of the lateral geniculate complex becomes apparent only when imaged using Ex-MSI (Fig. 3c). In Figs. 3d and 3e, the ion images display the distribution of PA (40:6) ([M-H]^-^, m/z 747.5) and SHexCer (42:2;O2) ([M-H]^-^, m/z 888.6) in the ventral posteromedial nucleus of the thalamus, highlighting the region’s complex and compact structure enriched with diverse lipid species. Despite the dilution and loss of lipids caused by sample expansion, we still achieved the detection of multiple species of lipid. Suppl. Fig. 3 displays the mass spectra of mouse brain tissues after expansion under positive and negative modes, respectively. There are 74 and 167 endogenous species that were found in positive and negative mode, respectively. Amongst, 125 of them were assigned as lipids with mass error within 5 ppm. The peak list with corresponding lipid assignments can be found in Suppl. Tables 2 and 3. To sum up, 19 lipid classes were detected. Suppl. Table 4 shows the list of CID fragments from selected lipids and their identities under positive and negative mode, respectively. This finding highlights the reliability of Ex-MSI for molecular imaging, demonstrating that the technique preserves sensitivity even under demanding conditions and offers high resolution.

**Fig. 3:**
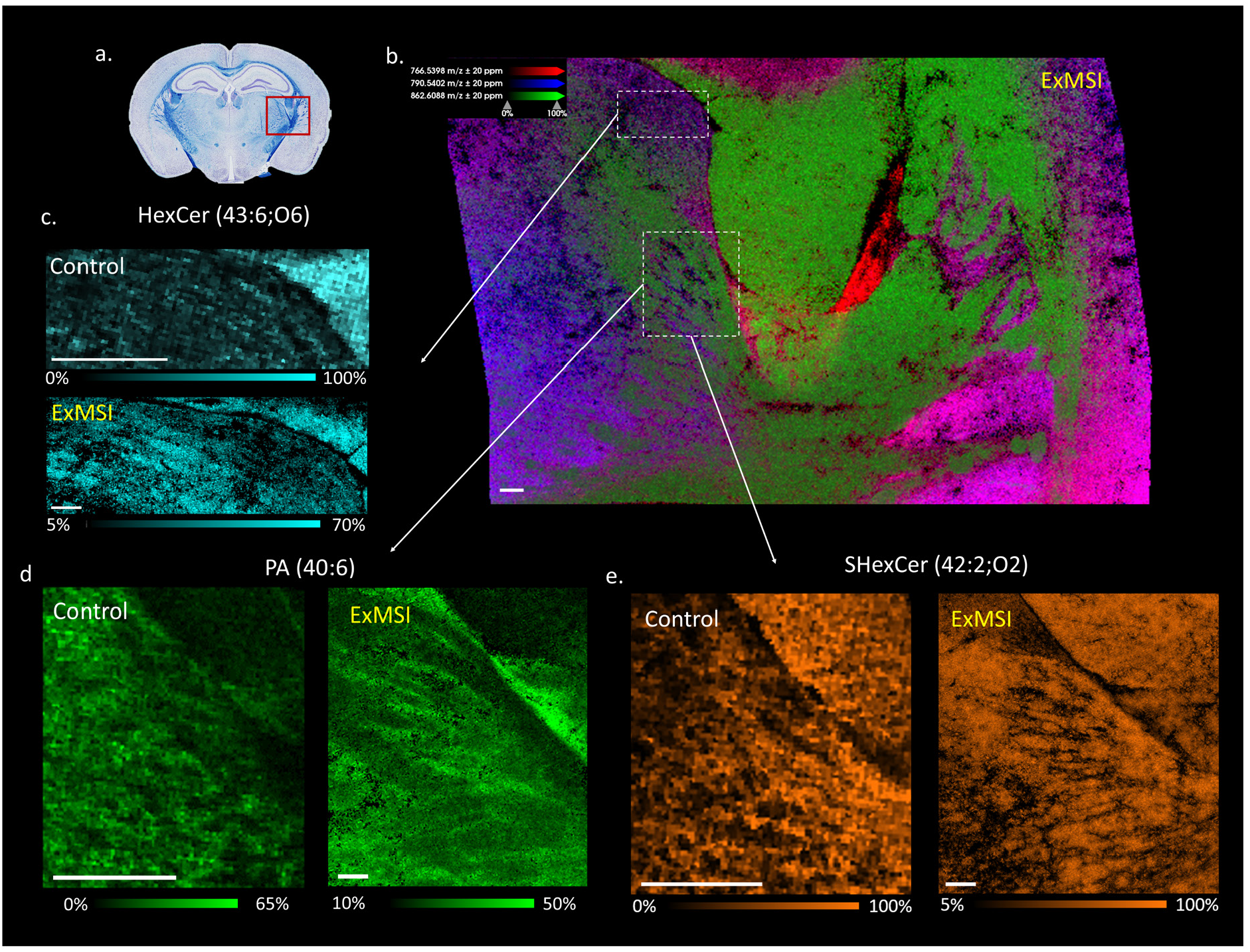
High-resolution lipid mapping in mouse brain cerebrum via Ex-MSI. Negative ion mode images were acquired using a 10 μm acquisition resolution, yielding an equivalent spatial resolution of 2.2 μm. (a) Coronal section of a mouse brain. Image courtesy from Bmonzon, Wikipedia, licensed under CC BY-SA 4.0. The red frame indicated the imaging area of (b). (b) Overlaid ion image of PE (38:4) ([M-H]^-^, m/z 766.5), PE (40:6) ([M-H]^-^, m/z 790.5), and SHexCer (40:1;O2) ([M-H]^-^, m/z 862.6). Panels (c-e) offer side-by-side comparisons between non-expanded and expanded ion images. (c) The fibrous structure of the dorsal part in the lateral geniculate complex could be observed with the ion images of HexCer (43:6;O6) ([M-H]^-^, m/z 878.6). (d, e) In the ventral posteromedial nucleus region of the thalamus, distribution of PA (40:6) ([M-H]^-^, m/z 747.5) and SHexCer (42:2;O2) ([M-H]^-^, m/z 888.6) were shown. Scale bar represents 300 μm.

Contrasting with fluorescence microscopy, which often requires complex labeling and staining to visualize specific compounds, MALDI MSI offers the distinctive ability to simultaneously detect a wide array of molecular entities in a single analysis. Importantly, MSI is particularly good at imaging small molecules with high specificity, a task that is often challenging in fluorescence-based microscopy. The introduction of Ex-MSI enhances this advantage by delivering exceptional spatial resolution without compromising molecular specificity.

## Conclusion

Expansion mass spectrometry imaging introduces a novel approach that bypasses the spatial resolution limitations of standard mass spectrometry instruments without the need for specialized instrumental modification. Ex-MSI notably enhances resolution by 4.5 times, achieving near 1 μm level. This technique provides an imaging tool for spatially detailed analysis of lipid metabolism in brain tissue. This technique represents a significant advancement in mass spectrometry imaging, a significant boost in resolution while harnessing the reliability of commercial instruments. This combination not only improves the quality of imaging but also ensures that the technique is robust and widely applicable, making it a reliable tool for advancing our understanding of brain functions, disorders, and the development of targeted treatments.

## Acknowledgments

This study was supported by the General Research Fund (12302122) of the Research Grants Council, Hong Kong Special Administrative Region, China, the Start-up Grant from Hong Kong Baptist University, and the SKLEBA Research Grant (SKLP_2021_P04). We thank Dr. Jie Liu, Southern University of Science and Technology, for her help in fixation and staining experiments in expansion microscopy. We thank Dr. Paul Tillberg, Janelia Research Campus, HHMI, for the consultant and discussion on expansion microscopy. We thank Dr. Fei Chen, Broad Institute of MIT and Harvard, for the consultant on expansion microscopy experiments. We thank Dr. Lin Zhu and Dr. Zhu Yang, Hong Kong Baptist University, for the discussion of relevant academic issues.

## Author Contributions

Conceptualization: Jianing Wang

Methodology: Yik Ling Winnie Hung, Jianing Wang

Validation: Chengyi Xie, Xin Diao, Ruxin Li

Formal Analysis: Yik Ling Winnie Hung

Investigation: Yik Ling Winnie Hung, Chengyi Xie, Xin Diao

Resources: Xiaoxiao Wang, Shulan Qiu, Jiacheng Fang

Visualization: Yik Ling Winnie Hung, Jianing Wang, Chengyi Xie

Writing - Original Draft: Yik Ling Winnie Hung, Jianing Wang

Writing - Review & Editing: Jianing Wang, Chengyi, Xie, Zongwei

Cai Supervision: Jianing Wang, Zongwei Cai

Project administration: Zongwei Cai

Funding acquisition: Jianing Wang, Zongwei Cai

## Notes

### Competing Interest Statement

The authors have declared no competing interest.

### Summary of Updates

In the updated version of our manuscript, we have made important improvements. We carefully reviewed our data and removed any that did not meet our quality standards. This step ensures the data's accuracy and relevance. We also corrected errors from the previous version, enhancing the clarity and correctness of our findings. By narrowing down our research focus, we have made our study more direct and precise. These revisions demonstrate our dedication to presenting clear and reliable research, and they enrich the manuscript's value to the scientific field.

