## Supplemental Information for "Expansion Strategy-Driven Micron-Level Resolution Mass Spectrometry Imaging of Lipids in Mouse Brain Tissue"

### Methods

#### Reagents and materials

Paraformaldehyde (4%) was bought from Beyotime Biotechnology (China). Glutaraldehyde, Triton X-100, sucrose, succinimidyl ester of 6−((acryloyl)amino)hexanoic acid (AcX), acrylamide (Acr), N,N'-methylenebisacrylamide (Bis), ammonium persulphate (APS), tetramethylethylenediamine (TEMED), sodium chloride, 2,5-dihydroxybenzoic acid (DHB), N-(1-naphthyl) ethylenediamine dihydrochloride (NEDC), 9-acridinamine (9AA), methanol (MeOH) were purchased from Sigma Aldrich (U.S.A.). Phosphate buffered saline (PBS), protease K, Tris-Cl (1 M, pH 8) were acquired from Thermo Fisher Scientific (U.S.A.). Sodium acrylate (SA) was obtained from Santa Cruz Biotechnology (U.S.A.). All reagents were used without further purification.

#### Animal

C57/BL mice of 8-week old were purchased from Laboratory Animal Service Centre, the Chinese University of Hong Kong. They were housed in a sterile and individually ventilated cages with 12-hr light/dark cycle at 22 °C and 45% relative humidity. All animal experiments were performed in accordance with Animals (Control of Experiments) Ordinance Chapter 340 approved by the Department of Health.

#### Preparation of tissue slices

Mice were anesthetized with 5% (v/v) isoflurane and rapidly decapitated. Brain was removed and post-fixed using 4% PFA with 0.1% glutaraldehyde in PBS overnight at 4 °C. They were washed with PBS for 24 hr and cryoprotected by incubating in 10% and 30% (w/v) sucrose solution at 4 °C until they sunk after 30 min and 24 hr, respectively. The brain was then frozen at -70 °C for storage. For expansion samples, 50-μm slices were sectioned using CryoStar$^{TM}$ NX70 cryostat (Thermo Fisher Scientific Inc., U.S.A.) at -20 °C. For non-expanded tissue, 10-μm slices were sectioned at the same cryostat conditions.

**Gelation**

Tissue slices were treated with 1xPBS at room temperature for 15 minutes. The slices were incubated with 0.1 mg/mL AcX in PBS overnight at room temperature, then washed with PBS twice for 15 minutes each. A monomer solution was prepared with 86 mg/mL sodium acrylate, 25 mg/mL acrylamide, 1.5 mg/mL N,N'-methylenebisacrylamide, and 117 mg/mL sodium chloride in PBS. The tissue slices were transferred to the gelation chamber, and 20 μL of the monomer solution was added for a 1-minute incubation. After drying the slices, the monomer solution was mixed with 10% (w/v) APS solution and 10% (v/v) TEMED solution in a 47:1:1 ratio. The resulting gelation solution was added to the chamber, covered with a Parafilm-wrapped glass, and the whole chamber was placed in a humid environment and incubated for 1 hour at 37 °C.

**Tissue digestion and expansion**

Excess gel around the tissue slice was trimmed off into an asymmetric shape. The gelation chamber was placed into a tailor-made ITO slide box filled with 8 units/mL proteinase K in a digesting solution of 50 mM Tris-Cl (pH 8), 0.5% Triton-X 100, 0.8 M NaCl, and digested at 60 °C for 3 hours in a humid chamber. The gel was detached and rinsed with 4 °C PBS for 10 minutes three times on ice, then expanded in 4 °C deionized water for 20 minutes three times on ice, until no further expansion occurred. The gel was quickly rinsed by 4 °C deionized water three times on ice, with liquid removed as much as possible, and The ITO slide box was placed in a drying chamber overnight.

**MALDI-MSI analysis**

For non-expanded tissue experiments, tissue slices were directly thaw-mounted onto Indium Tin Oxide (ITO) slides and dehydrated using a vacuum desiccator. Various matrices—DHB (20 mg/mL in MeOH), NEDC (5 mg/mL in MeOH) and 9AA (3 mg/mL in 95% MeOH) were applied using a custom-built matrix deposition device. Instrumental settings were as follows: matrix flow rate 10 µL/min, temperature 60 °C, XY-stage speed 10.98 cm/min, line-to-line distance of 1 mm, capillary voltage 5000 V, nitrogen sheath and auxiliary gas flow rate 70 and 5 psi, respectively. 10 cycles were sprayed. The MSI experiments were conducted using a timsTOF fleX instrument from Bruker Daltonics (Germany), equipped with a microGRID system. Mass spectra were acquired in both positive and negative modes with 50 – 450 laser shots. Lateral resolution was 5 – 20 µm in single mode and 50 µm in M5 small mode at a mass range of m/z 300 – 1300 and m/z 500 – 1100 for positive and negative mode, respectively. The laser power was set at 50 – 100 % and a repetition rate of 5000 Hz was used. ESI-L low-concentration tuning mix (Agilent Technologies, U.S.A.) was used for instrument calibration. Images were generated using SCiLS Lab MVS version 2022a Premium 3D (Bruker Daltonics, Germany). Feature selection was normalized to total ion count and executed with a mass tolerance of 20 ppm.

**Lipid identification**

Lipid species were assigned by comparing their measured accurate masses to the LIPID MAPS database, with a mass error threshold of less than 5 ppm. On-tissue MS/MS analysis was utilized for the further characterization of certain representative lipids, including PC, HexCer, PE, LPE, PA, and PI. Spectral interpretation was performed manually using two specialized tools: for negative ion mode, the Glycerophospholipid MS/MS Prediction and Product Ion Calculation tool available on LIPID MAPS was employed; for positive ion mode, the LipidBlast In-Silico MS/MS Database was utilized.

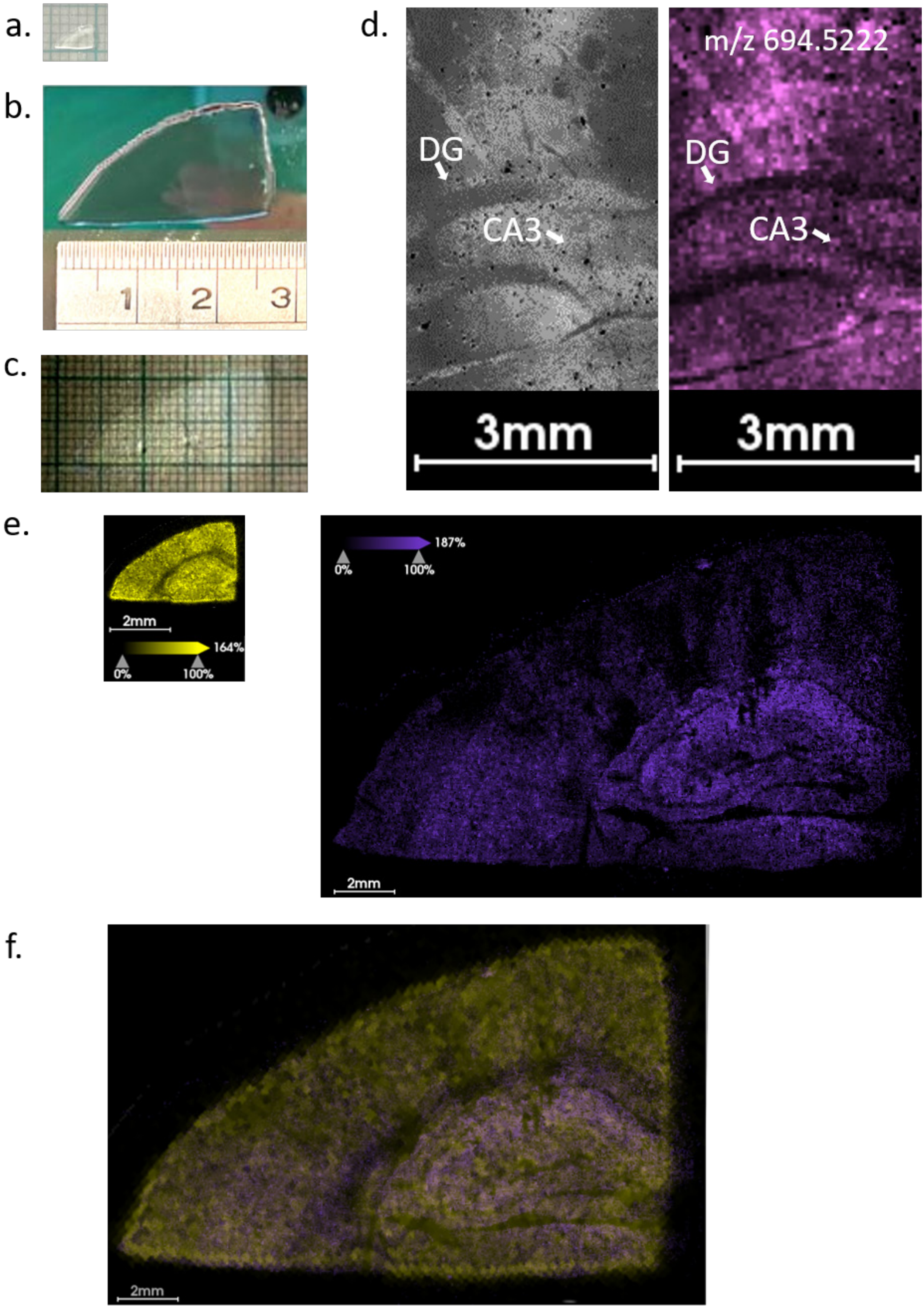
a.
b.
1
2
3
c.
d.
DG
CA3
3mm
m/z 694.5222
DG
CA3
3mm
e.
2mm
164%
0%
100%
187%
0%
100%
2mm
f.
2mm

Suppl. Fig. 1: Evaluation of Expansion Factor and Tissue Uniformity in Mouse Cerebrum Hippocampus Region.

(a) Pre-expansion hippocampus tissue section with a bottom width of 4.5 mm.

(b) Post-expansion tissue, displaying a bottom width of 26 mm.

(c) Tissue after drying, exhibiting a bottom width of 23 mm. The calculated expansion factor is 5.1.

(d) Comparative visualization of the hippocampus region via bright-field microscopy (left) and Mass Spectrometry Imaging (MSI) (right).

(e) Positive ionization mode MSI images showing ion at m/z 419.2 before (left) and after (right) expansion.

(f) Overlaid image combining data from (e).

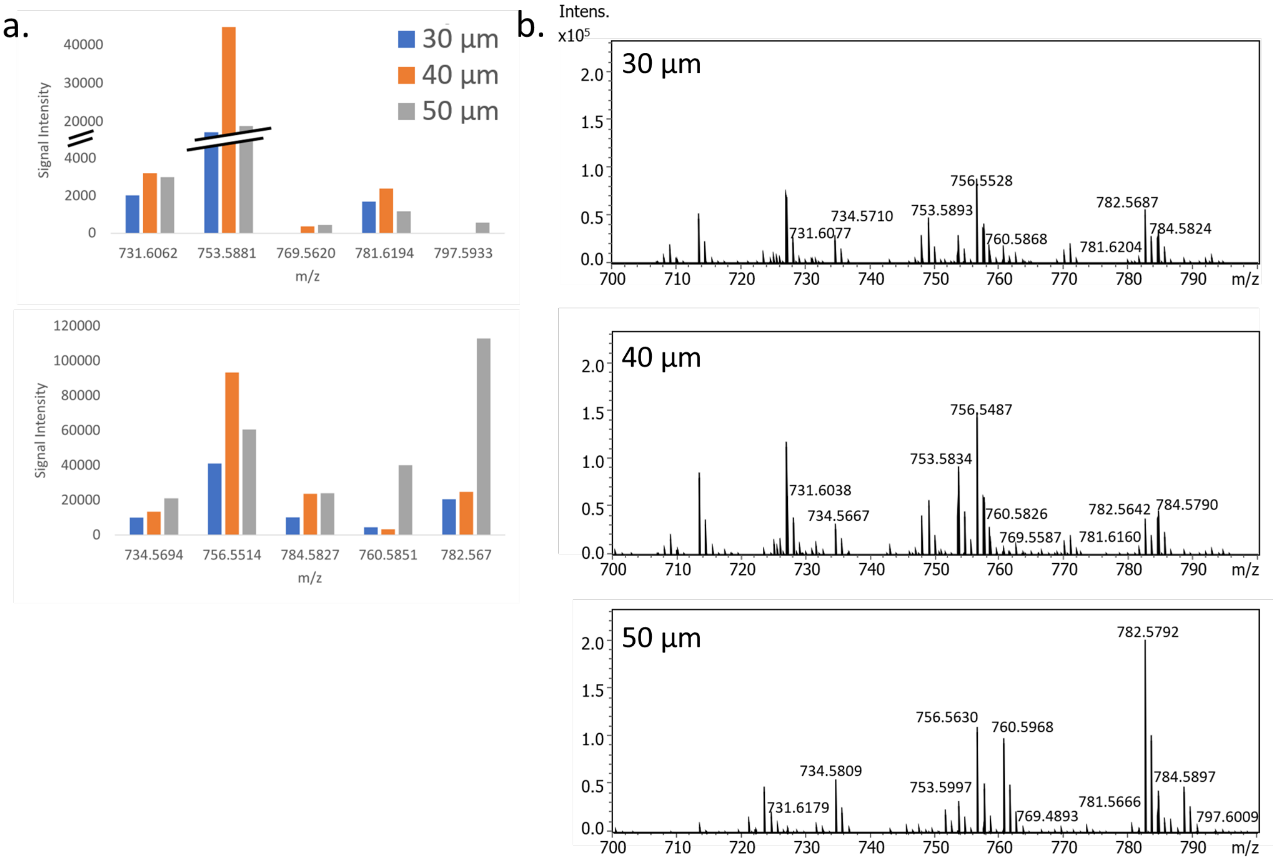


Suppl. Fig. 2: A representative signal comparison in lipid Ex-MSI across variable tissue thicknesses.

(a) Signal intensity comparison across 30, 40, and 50-µm tissue sections in lipid Ex-MSI.

(b) Representative negative ion mode mass spectra obtained from lipid Ex-MSI at different tissue section thicknesses (30, 40, and 50 µm) using NEDC as a matrix.

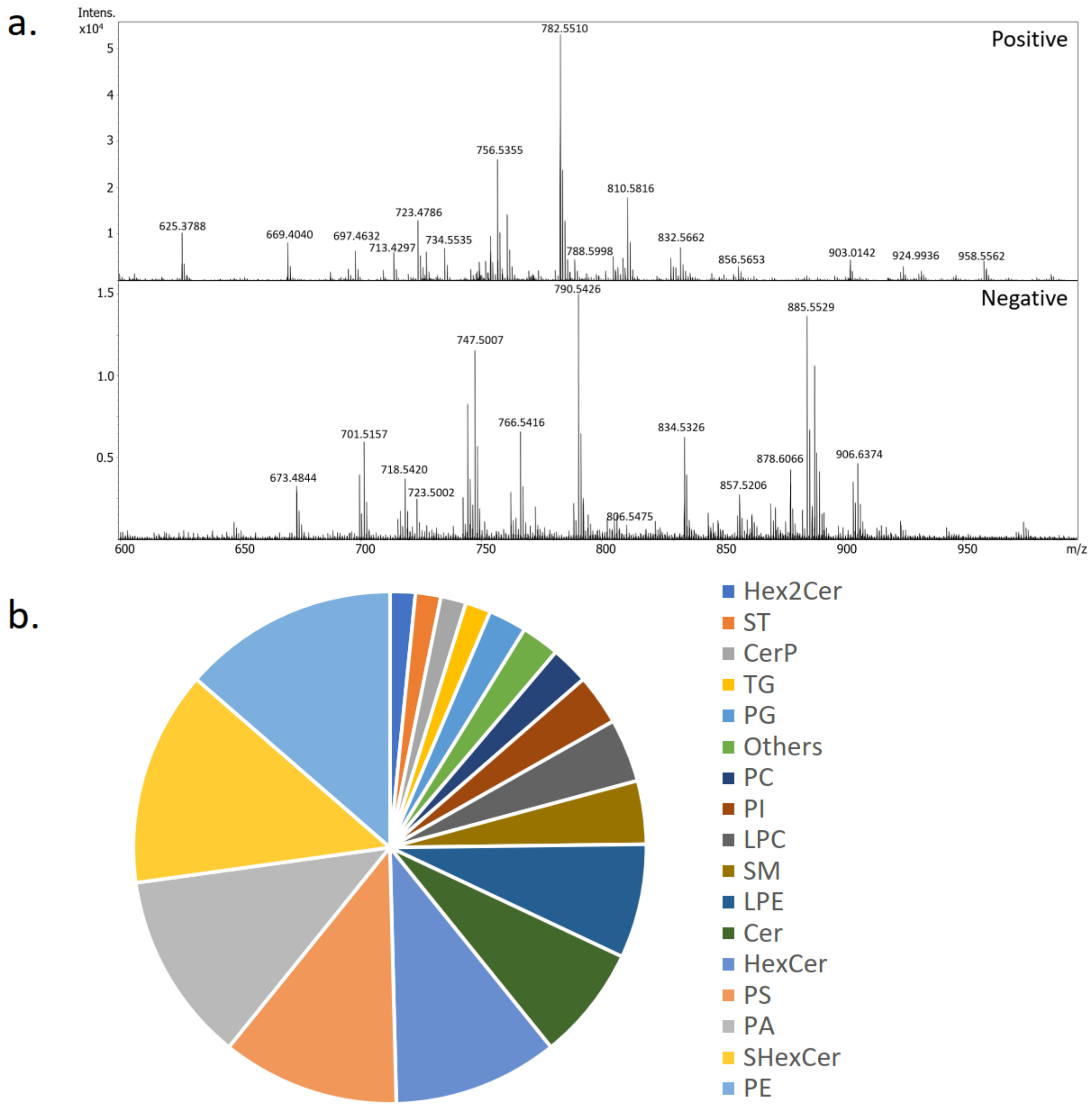


Suppl. Fig. 3: Characterization of Lipid Composition in Expanded Mouse Cerebrum Tissue Slices.

(a) Representative mass spectra obtained from expanded mouse cerebrum tissue slices, captured under both positive and negative ionization modes. A laser raster spot size of 10 μm yielded an equivalent lateral spatial resolution of 2.2 μm.

(b) Pie chart illustrating the relative distribution of various lipid classes detected in the expanded mouse cerebrum tissue.

Suppl. Table 1, Peak list of the selected lipids

| Theoretical m/z | Identity | Chemical Formula | Ion Type |
|---|---|---|---|
| 731.6061 | SM (36:1;O2) | C41H83N2O6P | $[M+H]^+$ |
| 734.5694 | PC (32:0) | C40H80NO8P | $[M+H]^+$ |
| 753.5881 | SM (36:1;O2) | C41H83N2O6PNa | $[M+Na]^+$ |
| 756.5514 | PC (32:0) | C40H80NO8PNa | $[M+Na]^+$ |
| 760.5851 | PC (34:1) | C42H82NO8P | $[M+H]^+$ |
| 769.5620 | SM (36:1;O2) | C41H83N2O6PK | $[M+K]^+$ |
| 781.6194 | SM (38:1;O2) | C43H87N2O6PNa | $[M+Na]^+$ |
| 782.5670 | PC (34:1) | C42H82NO8PNa | $[M+Na]^+$ |
| 784.5827 | PC (34:0) | C42H84NO8PNa | $[M+Na]^+$ |
| 797.5933 | SM (36:1;O2) | C43H87N2O6PK | $[M+K]^+$ |

Suppl. Table 2 List of lipid assignments within 5 ppm mass error for the ion species detected in expanded mouse cerebrum slice under positive mode. Laser pixel size was 20 μm.

| **Experimental m/z** | **Theoretical m/z** | **Error (ppm)** | **Identity** | **Chemical Formula** | **Ion Type** |
|---|---|---|---|---|---|
| 478.3312 | 478.3292 | 4.18 | LPC (O-16:2) | C24H48NO6P | $[M+H]^+$ |
| 518.3251 | 518.3241 | 1.93 | LPC (18:3) | C26H48NO7P | $[M+H]^+$ |
| 544.3417 | 544.3398 | 3.49 | LPC (20:4) | C28H50NO7P | $[M+H]^+$ |
| 674.3663 | 674.364 | 3.41 | PC (23:2;O3) | C31H58NO11PNa | $[M+Na]^+$ |
| 747.4926 | 747.4919 | 0.94 | SM (33:4;O6) | C38H71N2O10P | $[M+H]^+$ |
| 751.54 | 751.5385 | 2.00 | SM (37:6;O3) | C42H75N2O7P | $[M+H]^+$ |
| 753.5907 | 753.5881 | 3.45 | SM (36:1;O2) | C41H83N2O6PNa | $[M+Na]^+$ |
| 754.5521 | 754.5511 | 1.33 | LPC (O-33:2) | C41H82NO6PK | $[M+K]^+$ |
| 754.6087 | 754.6085 | 0.27 | LPC (O-34:1) | C42H86NO6PNa | $[M+Na]^+$ |
| 756.5543 | 756.5514 | 3.83 | PC (32:0) | C40H80NO8PNa | $[M+Na]^+$ |
| 760.5888 | 760.5851 | 4.86 | PC (34:1) | C42H82NO8P | $[M+H]^+$ |
| 769.4871 | 769.4893 | 2.86 | SM (34:3;O4) | C39H75N2O8PK | $[M+K]^+$ |
| 773.5181 | 773.5204 | 2.97 | SM (37:6;O3) | C42H75N2O7PNa | $[M+Na]^+$ |

Suppl. Table 3 List of lipid assignments within 5 ppm mass error for the ion species detected in expanded mouse cerebrum slice under negative mode. Laser pixel size was 10 μm.

| Experimental m/z | Theoretical m/z | Error (ppm) | Identity | Chemical Formula | Ion Type |
|---|---|---|---|---|---|
| 500.2776 | 500.2783 | 1.40 | LPE(20:4) | C25H44NO7P | [M-H]- |
| 502.2927 | 502.2939 | 2.39 | LPE (20:3) | C25H46NO7P | [M-H]- |
| 504.3084 | 504.3096 | 2.38 | LPE (20:2) | C25H48NO7P | [M-H]- |
| 506.3246 | 506.3252 | 1.19 | LPE (20:1) | C25H50NO7P | [M-H]- |
| 507.2622 | 507.26 | 4.34 | ST (21:2;O3;GlcA) | C27H40O9 | [M-H]- |
| 508.3387 | 508.3409 | 4.33 | LPE (20:0) | C25H52NO7P | [M-H]- |
| 524.2785 | 524.2783 | 0.38 | LPE (22:6) | C27H44NO7P | [M-H]- |
| 528.3098 | 528.3085 | 2.46 | LPE(22:4) | C27H48NO7P | [M-H]- |
| 536.254 | 536.255 | 1.86 | LPE (20:4) | C25H44NO7P | [M+Cl]- |
| 536.5409 | 536.5412 | 0.56 | Cer (35:0;O) | C35H71NO2 | [M-H]- |
| 540.2856 | 540.2862 | 1.19 | LPE (20:2) | C25H48NO7P | [M+Cl]- |
| 550.5197 | 550.5205 | 1.45 | Cer (35:1;O2) | C35H69NO3 | [M-H]- |
| 554.5144 | 554.5154 | 1.80 | Cer (34:0;O3) | C34H69NO4 | [M-H]- |
| 581.3091 | 581.3096 | 0.86 | PG (21:1;O) | C27H51O11P | [M-H]- |
| 599.3211 | 599.3202 | 1.50 | LPI (18:0) | C27H53O12P | [M-H]- |
| 600.575 | 600.5725 | 4.16 | Cer (40:3;O) | C40H75NO2 | [M-H]- |
| 616.5703 | 616.5674 | 4.70 | Cer (40:3;O2) | C40H75NO3 | [M-H]- |
| 630.531 | 630.5314 | 0.63 | Cer (36:0;O6) | C36H73NO7 | [M-H]- |
| 634.5793 | 634.578 | 2.05 | Cer (40:2;O3) | C40H77NO4 | [M-H]- |
| 647.4666 | 647.4657 | 1.39 | PA (32:0) | C35H69O8P | [M-H]- |
| 648.3802 | 648.3787 | 2.31 | ST (30:0;O8;T) | C32H59NO10S | [M-H]- |
| 650.3933 | 650.391 | 3.54 | HexCer (29:6;O4) | C35H57NO10 | [M-H]- |
| 652.4094 | 652.4066 | 4.29 | HexCer (29:5;O4) | C35H59NO10 | [M-H]- |
| 652.5888 | 652.5886 | 0.31 | Cer (40:1;O4) | C40H79NO5 | [M-H]- |
| 666.388 | 666.3907 | 4.05 | PE (28:2) | C33H62NO8P | [M+Cl]- |
| 673.4826 | 673.4814 | 1.78 | PA (34:1) | C37H71O8P | [M-H]- |

| | | | | | |
|---|---|---|---|---|---|
| 675.4959 | 675.497 | 1.63 | PA (34:0) | C37H73O8P | [M-H]- |
| 689.5002 | 689.4998 | 0.58 | TG (38:4;O2) | C41H70O8 | [M-H]- |
| 690.4973 | 690.495 | 3.33 | HexCer (34:5;O2) | C40H69NO8 | [M-H]- |
| 692.4991 | 692.5025 | 4.91 | CerP (40:5;O2) | C40H72NO6P | [M-H]- |
| 699.4973 | 699.497 | 0.43 | PA (36:2) | C39H73O8P | [M-H]- |
| 701.5113 | 701.5127 | 2.00 | PA (36:1) | C39H75O8P | [M-H]- |
| 706.5168 | 706.5181 | 1.84 | CerP (41:5;O2) | C41H74NO6P | [M-H]- |
| 712.5549 | 712.5522 | 3.79 | ACer (44:6;O4) | C44H75NO6 | [M-H]- |
| 715.5753 | 715.576 | 0.98 | CerPE (38:1;O2) | C40H81N2O6P | [M-H]- |
| 716.5241 | 716.5236 | 0.70 | PE (34:1) | C39H76NO8P | [M-H]- |
| 718.5404 | 718.5392 | 1.67 | PE (34:0) | C39H78NO8P | [M-H]- |
| 721.4816 | 721.4814 | 0.28 | PA (38:5) | C41H71O8P | [M-H]- |
| 723.4973 | 723.497 | 0.41 | PA (38:4) | C41H73O8P | [M-H]- |
| 725.5114 | 725.5127 | 1.79 | PA (38:3) | C41H75O8P | [M-H]- |
| 727.5275 | 727.5283 | 1.10 | PA (38:2) | C41H77O8P | [M-H]- |
| 728.5322 | 728.5318 | 0.55 | HexCer (34:2;O4) | C40H75NO10 | [M-H]- |
| 729.5419 | 729.544 | 2.88 | PA (38:1) | C41H79O8P | [M-H]- |
| 730.5457 | 730.5475 | 2.46 | HexCer (34:1;O4) | C40H77NO10 | [M-H]- |
| 742.54 | 742.5392 | 1.08 | PE (36:2) | C41H78NO8P | [M-H]- |
| 744.555 | 744.5549 | 0.13 | PE (36:1) | C41H80NO8P | [M-H]- |
| 747.4981 | 747.497 | 1.47 | PA (40:6) | C43H73O8P | [M-H]- |
| 748.4742 | 748.477 | 3.74 | PS (32:1;O) | C38H72NO11P | [M-H]- |
| 751.5267 | 751.5283 | 2.13 | PA (40:4) | C43H76O8P | [M-H]- |
| 759.4707 | 759.4737 | 3.95 | PA (38:4) | C41H73O8P | [M+Cl]- |
| 762.507 | 762.5079 | 1.18 | PE (38:6) | C43H74NO8P | [M-H]- |
| 764.5227 | 764.5236 | 1.18 | PE (38:5) | C43H76NO8P | [M-H]- |
| 766.5398 | 766.5392 | 0.78 | PE (38:4) | C43H78NO8P | [M-H]- |
| 770.5709 | 770.5705 | 0.52 | PE (38:2) | C43H82NO8P | [M-H]- |
| 772.5847 | 772.5862 | 1.94 | PE (38:1) | C43H84NO8P | [M-H]- |
| 776.5 | 776.5003 | 0.39 | PE (36:3) | C41H76NO8P | [M+Cl]- |

| | | | | | |
|---|---|---|---|---|---|
| 786.5266 | 786.5291 | 3.18 | PS (36:2) | C42H78NO10P | [M-H]- |
| 788.523 | 788.5236 | 0.76 | PE (40:7) | C45H76NO8P | [M-H]- |
| 788.5427 | 788.5447 | 2.54 | PS (36:1) | C42H80NO10P | [M-H]- |
| 790.5402 | 790.5392 | 1.26 | PE (40:6) | C45H78NO8P | [M-H]- |
| 794.5465 | 794.5472 | 0.88 | PE (37:1) | C42H82NO8P | [M+Cl]- |
| 794.5704 | 794.5705 | 0.13 | PE (40:4) | C45H82NO8P | [M-H]- |
| 798.649 | 798.6465 | 3.13 | HexCer (40:1;O3) | C46H89NO9 | [M-H]- |
| 802.4034 | 802.4053 | 2.37 | SHexCer (33:6;O5) | C39H65NO14S | [M-H]- |
| 802.5118 | 802.5145 | 3.36 | SHexCer (36:3;O2) | C42H77NO11S | [M-H]- |
| 804.5286 | 804.5316 | 3.73 | PE (38:3) | C43H80NO8P | [M+Cl]- |
| 806.5465 | 806.5458 | 0.87 | SHexCer (36:1;O2) | C42H81NO11S | [M-H]- |
| 810.5244 | 810.5221 | 2.84 | PS (38:4) | C44H78NO10P | [M-H]- |
| 816.5731 | 816.576 | 3.55 | PS (38:1) | C44H84NO10P | [M-H]- |
| 820.4159 | 820.4159 | 0.00 | SHexCer (33:5;O6) | C39H67NO15S | [M-H]- |
| 821.5448 | 821.5469 | 2.56 | PG (O-38:3) | C44H83O9P | [M+Cl]- |
| 822.5436 | 822.5421 | 1.82 | PS (O-37:2) | C43H82NO9P | [M+Cl]- |
| 826.6748 | 826.6778 | 3.63 | HexCer (42:1;O3) | C48H93NO9 | [M-H]- |
| 832.6067 | 832.6073 | 0.72 | PS (39:0) | C45H88NO10P | [M-H]- |
| 834.5281 | 834.5291 | 1.20 | PS (40:6) | C46H78NO10P | [M-H]- |
| 834.5781 | 834.5771 | 1.20 | SHexCer (38:1;O2) | C44H85NO11S | [M-H]- |
| 834.6199 | 834.6231 | 3.83 | HexCer (40:1;O3) | C46H89NO9 | [M+Cl]- |
| 835.5301 | 835.5283 | 2.15 | PA (47:11) | C50H77O8P | [M-H]- |
| 837.5462 | 837.5499 | 4.42 | PI (34:0) | C43H83O13P | [M-H]- |
| 838.556 | 838.5534 | 3.10 | Hex2Cer (30:0;O4) | C42H81NO15 | [M-H]- |
| 844.6407 | 844.6437 | 3.55 | PS(O-41:1) | C47H92NO9P | [M-H]- |
| 848.6368 | 848.6386 | 2.12 | PS (O-40:0;O) | C46H92NO10P | [M-H]- |
| 850.5739 | 850.572 | 2.23 | SHexCer | C44H85NO12S | [M-H]- |

| | | | | | |
|---|---|---|---|---|---|
| | | | (38:1;O3) | | |
| 851.5782 | 851.5808 | 3.05 | PG (42:5) | C48H85O10P | [M-H]- |
| 856.5103 | 856.5134 | 3.62 | PS (42:9) | C48H76NO10P | [M-H]- |
| 857.5172 | 857.5186 | 1.63 | PI (36:4) | C45H79O13P | [M-H]- |
| 860.5927 | 860.5927 | 0.00 | SHexCer (40:2;O2) | C46H87NO11S | [M-H]- |
| 860.6357 | 860.6386 | 3.37 | PS (41:0) | C47H92NO10P | [M-H]- |
| 862.6088 | 862.6084 | 0.46 | SHexCer (40:1;O2) | C46H89NO11S | [M-H]- |
| 862.6519 | 862.6544 | 2.90 | HexCer (42:1;O3) | C48H93NO9 | [M+Cl]- |
| 863.5625 | 863.5596 | 3.36 | PA (49:11) | C52H81O8P | [M-H]- |
| 864.6513 | 864.6488 | 2.89 | PE (45:4) | C50H92NO8P | [M-H]- |
| 870.4247 | 870.426 | 1.49 | Hex2Cer (28:4;O6) | C40H69NO17 | [M+Cl]- |
| 872.4246 | 872.4239 | 0.80 | SHexCer (34:4;O6) | C40H71NO15S | [M+Cl]- |
| 874.6084 | 874.6084 | 0.00 | SHexCer (41:2;O2) | C47H89NO11S | [M-H]- |
| 874.6434 | 874.6414 | 2.29 | HexCer (45:6;O4) | C51H89NO10 | [M-H]- |
| 876.5872 | 876.5876 | 0.46 | SHexCer (40:2;O3) | C46H87NO12S | [M-H]- |
| 876.6223 | 876.6206 | 1.94 | HexCer (44:6;O5) | C50H87NO11 | [M-H]- |
| 878.4388 | 878.4381 | 0.80 | PS (40:10;O) | C46H70NO11P | [M+Cl]- |
| 878.6013 | 878.6033 | 2.28 | SHexCer (40:1;O3) | C46H89NO12S | [M-H]- |
| 883.5308 | 883.5342 | 3.85 | PI (38:5) | C47H81O13P | [M-H]- |
| 885.5476 | 885.5499 | 2.60 | PI (38:4) | C47H83O13P | [M-H]- |
| 886.6064 | 886.6084 | 2.26 | SHexCer (42:3;O2) | C48H89NO11S | [M-H]- |
| 888.6213 | 888.624 | 3.04 | SHexCer (42:2;O2) | C48H91NO11S | [M-H]- |
| 890.6363 | 890.6363 | 0.00 | HexCer (45:6;O5) | C51H89NO11 | [M-H]- |
| 891.6388 | 891.6356 | 3.59 | TG (52:9;O3) | C55H88O9 | [M-H]- |
| 892.6202 | 892.6204 | 0.22 | PS (O-42:2) | C48H92NO9P | [M+Cl]- |

| | | | | | |
|---|---|---|---|---|---|
| 902.6015 | 902.6033 | 1.99 | SHexCer (42:3;O3) | $C_{48}H_{89}NO_{12}S$ | $[M-H]^-$ |
| 902.6376 | 902.6397 | 2.33 | SHexCer (43:2;O2) | $C_{49}H_{93}NO_{11}S$ | $[M-H]^-$ |
| 904.6165 | 904.6189 | 2.65 | SHexCer (42:2;O3) | $C_{48}H_{91}NO_{12}S$ | $[M-H]^-$ |
| 906.6315 | 906.6312 | 0.33 | HexCer (45:6;O6) | $C_{51}H_{89}NO_{12}$ | $[M-H]^-$ |
| 946.5804 | 946.5793 | 1.16 | IPC (40:1;O5) | $C_{46}H_{90}NO_{14}P$ | $[M+Cl]^-$ |

Suppl. Table 4. List of CID fragments from selected lipids and their identities under positive and negative modes

| Lipid | Experimental m/z | Identity | Theoretical m/z | Error (ppm) |
|---|---|---|---|---|
| PC (16:0/18:1) | 478.3261 | Neutral loss of sn-2 fatty acyl chain (18:1) from $[M+H]^+$ | 478.3292 | 6.48 |
| | 504.3425 | Neutral loss of sn-1 fatty acyl chain (16:0) from $[M+H]^+$ | 504.3449 | 4.76 |
| | 577.5163 | Neutral loss of phosphocholine from $[M+H]^+$ | 577.5190 | 4.68 |
| | 599.4980 | Neutral loss of phosphocholine from $[M+Na]^+$ | 599.5010 | 5.00 |
| | 723.4909 | Neutral loss of choline from $[M+Na]^+$ | 723.4935 | 3.59 |
| | 782.5643 | PC (16:0/18:1) precursor ion, $[M+Na]^+$ | 782.5670 | 3.45 |
| PC (16:0/16:0) | 478.3277 | Neutral loss of sn-1/2 fatty acyl chain (16:0) from $[M+H]^+$ | 478.3292 | 3.14 |
| | 500.3115 | Neutral loss of sn-1/2 fatty acyl chain (16:0) from $[M+Na]^+$ | 500.3111 | 0.80 |
| | 551.5006 | Neutral loss of phosphocholine from $[M+H]^+$ | 551.5034 | 5.08 |
| | 573.4804 | Neutral loss of phosphocholine from $[M+Na]^+$ | 573.4853 | 8.54 |
| | 697.4711 | Neutral loss of choline from $[M+Na]^+$ | 697.4779 | 9.75 |
| | 756.5441 | PC (16:0/16:0) precursor ion, $[M+Na]^+$ | 756.5514 | 9.65 |
| HexCer (18:1/24:0;2OH) | 484.3252 | Loss of sn-2 fatty acyl chain (24:0;2OH) from $[M+Na]^+$ | 484.3245 | 1.45 |
| | 649.6410 | Neutral loss of galactose head group from $[M+H]^+$ | 649.6367 | 6.62 |
| | 671.6182 | Neutral loss of galactose head | 671.6187 | 0.74 |

| | | | | |
|---|---|---|---|---|
| | | group from $[M+Na]^+$ | | |
| | 850.6733 | HexCer (18:1/24:0;2OH) precursor ion, $[M+Na]^+$ | 850.6743 | 1.18 |
| PC (18:0/18:1) | 526.3275 | Neutral loss of sn-1 fatty acyl chain (18:0) from $[M+Na]^+$ | 526.3268 | 1.33 |
| | 528.3490 | Neutral loss of sn-2 fatty acyl chain (18:1) from $[M+Na]^+$ | 528.3424 | 12.49 |
| | 605.5496 | Neutral loss of phosphocholine from $[M+H]^+$ | 605.5503 | 1.16 |
| | 627.5324 | Neutral loss of phosphocholine from $[M+Na]^+$ | 627.5323 | 0.16 |
| | 751.5254 | Neutral loss of choline from $[M+Na]^+$ | 751.5248 | 0.80 |
| | 810.5977 | PC (18:0/18:1) precursor ion, $[M+Na]^+$ | 810.5983 | 0.74 |
| PC (16:0/22:6) | 478.3287 | Neutral loss of sn-2 fatty acyl chain (22:6) from $[M+H]^+$ | 478.3292 | 1.05 |
| | 500.3091 | Neutral loss of sn-2 fatty acyl chain (22:6) from $[M+Na]^+$ | 500.3111 | 4.00 |
| | 550.3270 | Neutral loss of sn-1 fatty acyl chain (16:0) from $[M+H]^+$ | 550.3292 | 4.00 |
| | 572.3109 | Neutral loss of sn-1 fatty acyl chain (16:0) from $[M+Na]^+$ | 572.3111 | 0.35 |
| | 645.4856 | Neutral loss of phosphocholine from $[M+Na]^+$ | 645.4853 | 0.46 |
| | 769.4781 | Neutral loss of choline from $[M+Na]^+$ | 769.4779 | 0.26 |
| | 828.5512 | PC (16:0/22:6) precursor ion, $[M+Na]^+$ | 828.5514 | 0.24 |

| **Lipid** | Experimental m/z | Identity | Theoretical m/z | Error (ppm) |
|---|---|---|---|---|
| PI (18:0/20:4) | 283.2633 | sn1 RCOO- ion | 283.2643 | 3.53 |

| | | | | |
|---|---|---|---|---|
| | 303.2300 | sn2 RCOO- ion | 303.2319 | 6.27 |
| | 419.2557 | Neutral loss of sn-2 fatty acyl chain (20:4) and myo-inositol head group | 419.2568 | 2.62 |
| | 581.3111 | Loss of sn-2 fatty acyl chain (20:4) | 581.3096 | 2.58 |
| | 599.3167 | Loss of sn-2 fatty acyl chain (20:4) as ketene | 599.3202 | 5.84 |
| | 601.2759 | Loss of sn-1 fatty acyl chain (18:0) | 601.2783 | 3.99 |
| | 619.2871 | Loss of sn-1 fatty acyl chain (18:0) as ketene | 619.2889 | 2.91 |
| | 885.5488 | PI (18:0/20:4) precursor ion | 885.5499 | 1.24 |
| PE (18:0/18:1) | 283.2633 | sn1 RCOO- ion | 283.2643 | 3.53 |
| | 281.2497 | sn2 RCOO- ion | 281.2475 | 7.82 |
| | 419.2556 | Neutral loss of sn-2 fatty acyl chain (18:1) and ethanolamine head group | 419.2568 | 2.86 |
| | 437.2666 | Loss of sn-2 fatty acyl chain (18:1) as ketene and neutral loss of ethanolamine head group | 437.2674 | 1.83 |
| | 460.2819 | Neutral loss of sn-1 fatty acyl chain (18:0) | 460.2833 | 3.04 |
| | 462.2968 | Neutral loss of sn-2 fatty acyl chain (18:1) | 462.2990 | 4.76 |
| | 478.2956 | Loss of sn-1 fatty acyl chain (18:0) as ketene | 478.2939 | 3.55 |
| | 480.3081 | Loss of sn-2 fatty acyl chain (18:1) as ketene | 480.3096 | 3.12 |
| | 744.5538 | PE (18:0/18:1) precursor ion | 744.5549 | 1.48 |
| PA (18:0/18:1) | 283.2633 | sn1 RCOO- ion | 283.2643 | 3.53 |
| | 281.2482 | sn2 RCOO- ion | 281.2475 | 2.49 |
| | 417.2395 | Neutral loss of sn-1 fatty acyl chain (18:0) | 417.2412 | 4.07 |

| | | | | |
|---|---|---|---|---|
| | 419.2544 | Neutral loss of sn-2 fatty acyl chain (18:1) | 419.2568 | 5.72 |
| | 435.2501 | Loss of sn-1 fatty acyl chain (18:0) as ketene | 435.2517 | 3.68 |
| | 437.2647 | Loss of sn-2 fatty acyl chain (18:1) as ketene | 437.2674 | 6.17 |
| | 701.5121 | PA (18:0/18:1) precursor ion | 701.5127 | 0.86 |
| LPE (22:6) | 283.2429 | Loss of CO2 from sn-1 fatty acyl chain (22:6) | 283.2431 | 0.71 |
| | 327.2314 | sn-1 fatty acyl chain (22:6) | 327.2330 | 4.89 |
| | 524.2784 | LPE (22:6) precursor ion | 524.2783 | 0.19 |
| PE (16:0/18:0) | 255.2339 | sn1 RCOO- ion | 255.2319 | 7.84 |
| | 283.2660 | sn2 RCOO- ion | 283.2643 | 6.00 |
| | 419.2528 | Neutral loss of sn-1 fatty acyl chain (16:0) and ethanolamine head group | 419.2568 | 9.54 |
| | 437.2654 | Loss of sn-1 fatty acyl chain (16:0) as ketene and neutral loss of ethanolamine head group | 437.2674 | 4.57 |
| | 452.2763 | Loss of sn-2 fatty acyl chain (18:0) as ketene | 452.2783 | 4.42 |
| | 462.2934 | Neutral loss of sn-1 fatty acyl chain (16:0) | 462.2990 | 12.11 |
| | 480.3081 | Loss of sn-1 fatty acyl chain (16:0) as ketene | 480.3096 | 3.12 |
| | 718.5376 | PE (16:0/18:0) precursor ion | 718.5392 | 2.23 |
| PE (18:0/20:4) | 283.2713 | sn1 RCOO- ion | 283.2643 | 24.7 |
| | 303.2341 | sn2 RCOO- ion | 303.2319 | 7.26 |
| | 462.2934 | Neutral loss of sn-2 fatty acyl chain | 462.2990 | 12.11 |

| | | | | |
|---|---|---|---|---|
| | | (20:4) | | |
| | 480.3084 | Loss of sn-2 fatty acyl chain (20:4) as ketene | 480.3096 | 2.50 |
| | 500.2783 | Loss of sn-1 fatty acyl chain (18:0) as ketene | 500.2783 | 0.00 |
| | 766.5391 | PE (18:0/20:4) precursor ion | 766.5392 | 0.13 |
| PE (18:0/22:6) | 283.2661 | sn1 RCOO- ion | 283.2643 | 6.35 |
| | 327.2308 | sn2 RCOO- ion | 327.2330 | 6.72 |
| | 462.2968 | Neutral loss of sn-2 fatty acyl chain (22:6) | 462.2990 | 4.76 |
| | 480.3081 | Loss of sn-2 fatty acyl chain (22:6) as ketene | 480.3096 | 3.12 |
| | 506.2636 | Neutral loss of sn-1 fatty acyl chain (18:0) | 506.2677 | 8.10 |
| | 790.5371 | PE (18:0/22:6) precursor ion | 790.5392 | 2.66 |
